# Improved data-driven collective variables for biased sampling through iteration on biased data

**DOI:** 10.1101/2025.03.25.644418

**Authors:** Subarna Sasmal, Martin McCullagh, Glen M. Hocky

## Abstract

Our ability to efficiently sample conformational transitions between two known states of a biomolecule using collective variable (CV) based sampling depends strongly on the choice of the CV. We previously reported a data-driven approach to clustering biomolecular configurations with a probabilistic clustering model termed ShapeGMM. ShapeGMM is a Gaussian Mixture Model in cartesian coordinates, with means and covariances in each cluster representing the harmonic approximation to the conformational ensemble around a metastable state. We subsequently showed that Linear Discriminant Analysis on positions (posLDA) is a good reaction coordinate to characterize the transition between two of these states, and moreover can be biased to produce transitions between the states using Metadynamics-like approaches. However, the quality of these LDA coordinates depends on the amount of data used to characterize the states, and here we demonstrate the ability to systematically improve them using enhanced sampling data. Specifically, we demonstrate that improved CVs for sampling can be generated by iteratively performing biased sampling along a posLDA coordinate and then generating a new shapeGMM model from biased data in the previous iteration. The new coordinates derived from our iterative approach show a substantial improvement in being able to induce transitions between metastable states, and to converge a free energy surface.

## 1 Introduction

Molecular dynamics (MD) is a powerful approach for studying complex biochemical processes. ^1^ However, many critical events, such as protein folding and allosteric regulation of enzymes, occur on timescales that are often inaccessible to conventional MD due to the so-called rare event problem. ^1,2^ In these cases, the system becomes trapped in an initial metastable state, unable to overcome high free energy barriers that separate different regions of the free energy landscape. This limitation is particularly pronounced in large systems with many degrees of freedom, where fully sampling all relevant states is nearly impossible, even with very long MD simulations.

Over the years, numerous enhanced sampling techniques have been developed to alleviate this challenge by facilitating more frequent transitions between different states of a system. ^3,4^ One prominent class of these methods relies on collective variables (CVs), where an external bias is applied as a function of carefully chosen CVs. An ideal CV is thought to capture the slowest modes of motion responsible for significant conformational changes in macromolecules. By applying a bias to enhance fluctuations in one or several CVs, these methods encourage the system to explore low-probability regions of the free energy surface. However, the effectiveness of CV-based enhanced sampling techniques depends heavily on the choice of CV, which can be particularly challenging for complex systems.

There are numerous methods designed to identify “optimal” CVs for a given system, each with its own strengths and limitations. Some approaches employ simple linear dimensionality reduction techniques, while others leverage machine learning (ML) and deep learning algorithms to construct sophisticated nonlinear coordinates. ^5–12^ Interestingly, most of these methods rely on training with initial, under-sampled MD simulation data, which often lacks sufficient information about different metastable states and their transitions. The effectiveness of these approaches is inherently dependent on the quality of the sampling used for training. The resulting CV, obtained from this limited dataset, serves as a fixed reaction coordinate that is subsequently biased in enhanced sampling simulations to achieve a well-converged free energy surface (FES).

Recent studies have introduced a different strategy for identifying optimal reaction coordinates. These approaches employ an iterative scheme that refines the initially defined CV on-the-fly by leveraging reweighted data from successive biased simulations. ^13–22^ Unlike traditional methods where the CV remains fixed, this adaptive process continuously improves the coordinate as it is trained on progressively better-sampled data from each iteration of the biased simulations. This iterative refinement enhances the accuracy and efficiency of the CV, leading to a more reliable exploration of the free energy landscape. While these methods are highly efficient, they are computationally expensive. The resulting coordinates often lack clear physical interpretability and are sensitive to hyperparameters and neural network architecture, requiring careful tuning for optimal performance.

Here, we present an iterative scheme to improve our previously reported posLDA approach. ^9^ The iterative process starts by creating an initial CV using data from short, unbiased MD simulations. Enhanced sampling is then performed along this coordinate, and the free energy surface is assessed for convergence. If convergence is not achieved, biased samples are reweighted and clustered using our frame-weighted ShapeGMM ^23^ method to further refine the coordinate. This process repeats until the free energy surface or another relevant observable converges, providing an optimal reaction coordinate for efficient sampling. We have applied this protocol on two systems of increasing complexity-a nine residue peptide (Aib)_9_ and a 35-amino acid fast-folding Nle/Nle mutant Villin headpiece (also known as HP35). In both cases, iteration improves both the stability (meaning more aggressive biasing parameters can be used) and the sampling ability of the CV. This approach is implemented in tools that we have made available within an updated ShapeGMMTorch python package, ^24^ with biased simulations available in a number of MD simulation packages via our `sizeshape` PLUMED module. ^9,25^

## 2 Theory and Methods

### 2.1 Iteration Process

The iterative scheme employed in this work combines our previous three procedures ^9,23,26^ in a straightforward yet effective approach, as illustrated in the flowchart (Fig. 1). Our focus is on identifying a reaction coordi-nate that connects two specific states of a given system.

**Figure 1.**
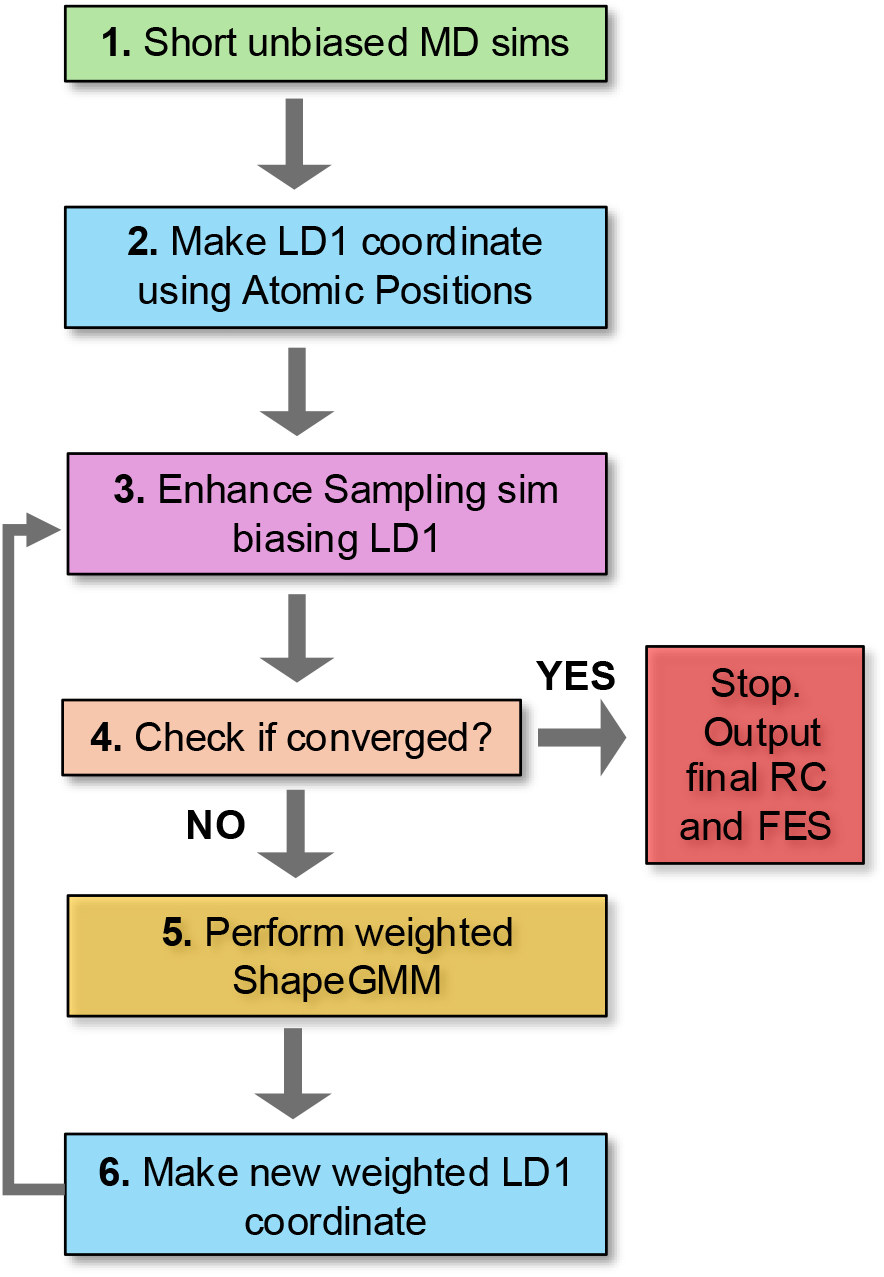
Workflow for iteration scheme. There are 6 steps: **1**. Short unbiased MD simulations are performed starting from two states of interest; **2**. use LDA method to make LD1 coordinate; **3**. perform enhanced sampling simulation, biasing LD1 coordinate; **4**. check the convergence of FES within a given threshold; if not converged, **5**. apply frame-weighted ShapeGMM on biased data to identify new states; **6**. apply frame-weighted LDA between two newly identified states to generate a new weighted LD1 coordinate. Repeat steps 3 to 6 until the simulation converges.

The process begins with two short unbiased MD simulations initiated from each state (or data from a long unbiased MD trajectory can be used if both states of interest are sufficiently well represented). From the resulting labeled MD simulation data, an initial linear discriminant analysis (LDA) coordinate is constructed. In the next step, an enhanced sampling technique such as Metadynamics (MetaD) or the On-the-fly Probability Enhanced Sampling variant (OPES-MetaD) ^27,28^ is used, applying a bias along the LDA coordinate within an MD simulation (see Sec. 2.4). Following this, the convergence of the free energy (FE) is assessed. If the FE surface has not yet converged, the workflow proceeds to the next stage, where the frame-weighted ShapeGMM method is used to cluster the biased samples. This method accounts for the non-uniform weights associated with each biased sample, effectively reweighting them to provide an unbiased estimate of the clusters. Subsequently, the newly identified clusters are analyzed to determine which two new clusters best correspond to the original state definitions

A new, weighted LDA coordinate is then computed using reweighted samples from these clusters. The process iterates through steps 3 to 6 until the free energy surface (FES) or another relevant physical observable converges within a predefined threshold. The reaction coordinate obtained at the end of this iterative process serves as an optimized bias coordinate, enhancing the efficiency of FES sampling when employed in enhanced sampling simulations. In practice, it may be difficult to assess convergence of the FES given finite time of sampling available, and so here we also focus on the efficiency of exploration, namely how frequently the biased CV and other physically intuitive CVs of interest transit between the values for the two target states.

### 2.2 Weighted ShapeGMM

In shapeGMM, a particular configuration of a macromolecule is represented by a particle position matrix, ***x***_*i*_, of order *N* × 3, where *N* is the number of particles being considered for clustering. To account for translational and rotational invariance, the proper feature for clustering purposes is an equivalence class,

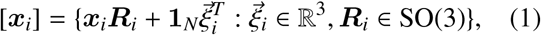

where 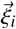 is a translation in R^3^, ℝ_*i*_ is a rotation ℝ^3^ → ℝ^3^, and **1**_*N*_ is the *N* × 1 vector of ones. [***x***_*i*_] is thus the set of all rigid body transformations, or orbit, of ***x***_*i*_.

The shapeGMM probability density is a Gaussian mixture given by

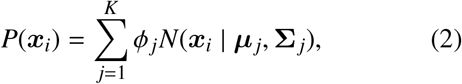

where the sum is over the *K* Gaussian mixture components, ϕ _*j*_ is the weight of component *j*, and *N*(***x***_*i*_ | ***µ*** _*j*_, **Σ** _*j*_) is a normalized multivariate Gaussian given by

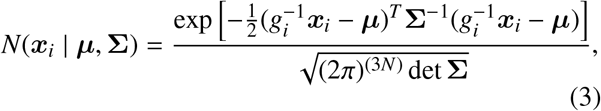

where ***µ*** is the mean structure, **Σ** is the covariance, and 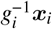 is the element of the equivalence class, [***x***_*i*_], that minimizes the squared Mahalanobis distance in the argument of the exponent. Determining the proper transformation, *g*_*i*_, is achieved by translating all frames to the origin and then determining an optimal rotation matrix. Cartesian and quaternion-based algorithms for determining optimal rotation matrices are known for two forms of the covariance were considered **Σ** ∝ ***I***_3*N*_ ^29,30^ or **Σ** = **Σ**_*N*_ ⊗***I***_3_, ^31,32^ where **Σ**_*N*_ is the *N* × *N* covariance matrix and ⊗ denotes a Kronecker product. In this manuscript, we employ only the more general Kronecker product covariance.

While using input data from an enhanced sampling simulation, we take non-uniform frame weights into account by performing weighted averages in the Expectation Maximization estimate of model parameters 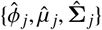. Considering a normalized set of frame weights, {*w*_*i*_} where 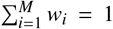 for *M* frames, their contribution to the probability can be accounted for by weighting the estimate of the posterior distribution of latent variables:

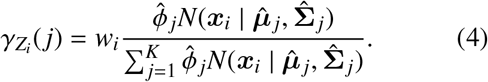

The frame weight will propagate to the estimate of component weights, means, and covariances in the Maximization step through 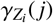:

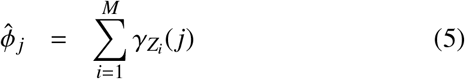

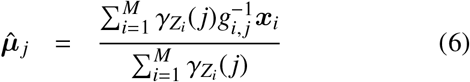

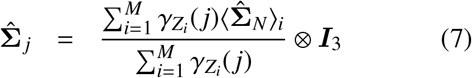

Additionally, the log likelihood per frame is computed as a weighted average

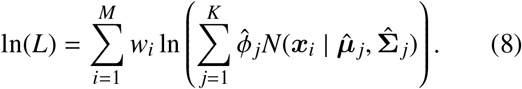

### 2.3 Frame-weighted LDA (wLDA)

LDA is a supervised classification technique that reduces dimensionality of the data by means of projection into a lower dimensional space. ^33^ It does so by simultaneously maximizing the between class scatter matrix and minimizing the within class scatter matrix. In our prior work, we have demonstrated that application of LDA on aligned particle positions produce a good one dimensional reaction coordinate that best separates two states. In Frame-weighted LDA approach, one can use input data obtained from enhanced sampling by incorporating nonuniform weights of the samples to account for relative probabilities of different classes. To do so, we must include weights corresponding to each configuration in the while computing the scatter matrices. For *K* different clusters, this is achieved by first computing the weighted within-cluster scatter matrix,

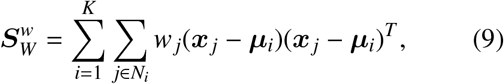

and the between-cluster scatter matrix,

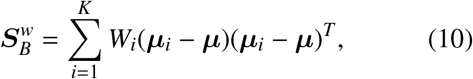

where 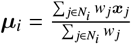 is the weighted mean of cluster *i*, {*w*_*j*_} are the normalized weights of individual samples such that 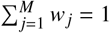 and *N*_*i*_ is the number of samples that belong to cluster *i*. 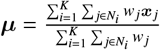 is the overall weighted global mean and 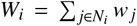 is the total weight of samples in cluster *i*. The simultaneous minimization of within-cluster scatter and maximization of between cluster scatter can be achieved by finding the transformation *G* that maximizes the quantity

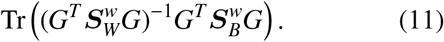

This maximization can be achieved through an eigenvalue/eigenvector decomposition but such a procedure is only applicable when 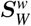 is non-singular. The LDA method was reformulated in terms of the generalized singular value decomposition (SVD) ^34^ extending the applicability of the method to singular 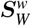 matrices such as those encountered when using particle positions.

We have implemented this modified approach in a WeightedLDA python package. ^35^ The result of an wLDA procedure on two labeled states will be a vector, ***v***, of coefficients that best separate the two states. These coefficients can be used directly for sampling within our PLUMED `sizeshape` module.

### 2.4 Enhanced sampling with LDA coordinates

The LDA coordinate used here is a dot product of the vector ***v*** with the atomic coordinates ***x*** − ***µ***, and it is given by, ^9^s

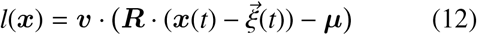

To compute the value of the LDA coordinate *l* on the fly, we first translate 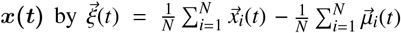, the difference in the geometric mean of the current frame and that of the reference configuration. Then, we compute ***R***(*t*), the rotation matrix which minimizes the Mahalanobis difference between 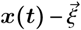 and ***µ***, for a given **Σ**, as described in Ref. 26.

Enhanced sampling simulations on LDA coordinates were performed using Well-tempered Metadynamics (WT-MetaD), and On the Fly Probability Enhanced Sampling-Metadynamics (OPES-MetaD) as implemented in PLUMED. ^25,28,36,37^

WT-MetaD works by adding a bias formed from a history dependent sum of progressively shrinking Gaussian hills. ^27,38^ The bias at time *t* for CV value *s*_*i*_ is given by the expression

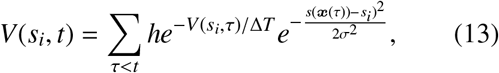

where *h* is the initial hill height, σ sets the width of the Gaussians, and Δ*T* is an effective sampling temperature for the CVs. Rather than setting Δ*T* , one typically chooses the bias factor γ = (*T* + Δ*T* )/*T* , which sets the smoothness of the sampled distribution. ^27,38^ Asymptotically, a free energy surface (FES) can be estimated from the applied bias by 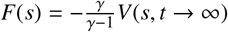 ^27,39^ or using a reweighting scheme. ^27,40^

OPES-MetaD applies a bias that is based on a kernel density estimate of the probability distribution over the whole space, which is iteratively updated. ^28,37^ The bias at time *t* for CV value *s*_*i*_ takes the form

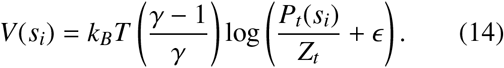

Here in the prefactor, *T* is the temperature, *k*_*B*_ is Boltzmann’s constant, and γ is the bias factor. *P*_*t*_(*s*) is the current estimate of the probability distribution, *Z*_*t*_ is a normalization factor that comes from integrating over sampled *s* space. Finally, 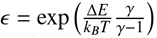 is a regularization constant that ensures the maximum bias that can be applied is Δ*E*.

As in WT-MetaD, *F*(*s*) can be directly estimated from *V*(*s*) by 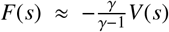 or through a reweighting scheme. ^37^ Details of the sampling parameters used for each system are given in Sec. 5.

## 3 Results and Discussion

### 3.1 Performing iterations on (Aib)_9_

(Aib)_9_ is a nine residue peptide formed from the achiral α-aminoisobutyryl that exhibits two well defined metastable states: left- and right-handed α-helices. Due to the symmetry inherent in a helix made of achiral building blocks, both states must have equal statistical likelihood. In work by us and others, ^9,15,41^ this symmetry was leveraged to benchmark sampling and clustering methods. Here, we applied the proposed iterative scheme on this system to assess the improvement of the reaction coordinate over successive iterations.

The iterative process begins with an initial linear discriminant (LD1) coordinate derived from two short molecular dynamics (MD) simulations starting from the left- and right-handed states. This is followed by a WT-MetaD simulation biasing this coordinate (for details, see Ref. 9). The LD coordinate exhibited several transitions between extreme values representing the left and right helical states (Fig. 2, top) within 500 ns. We subsequently performed three more iterations, with each WT-MetaD simulation also running for 500ns (Fig. 2).

**Figure 2.**
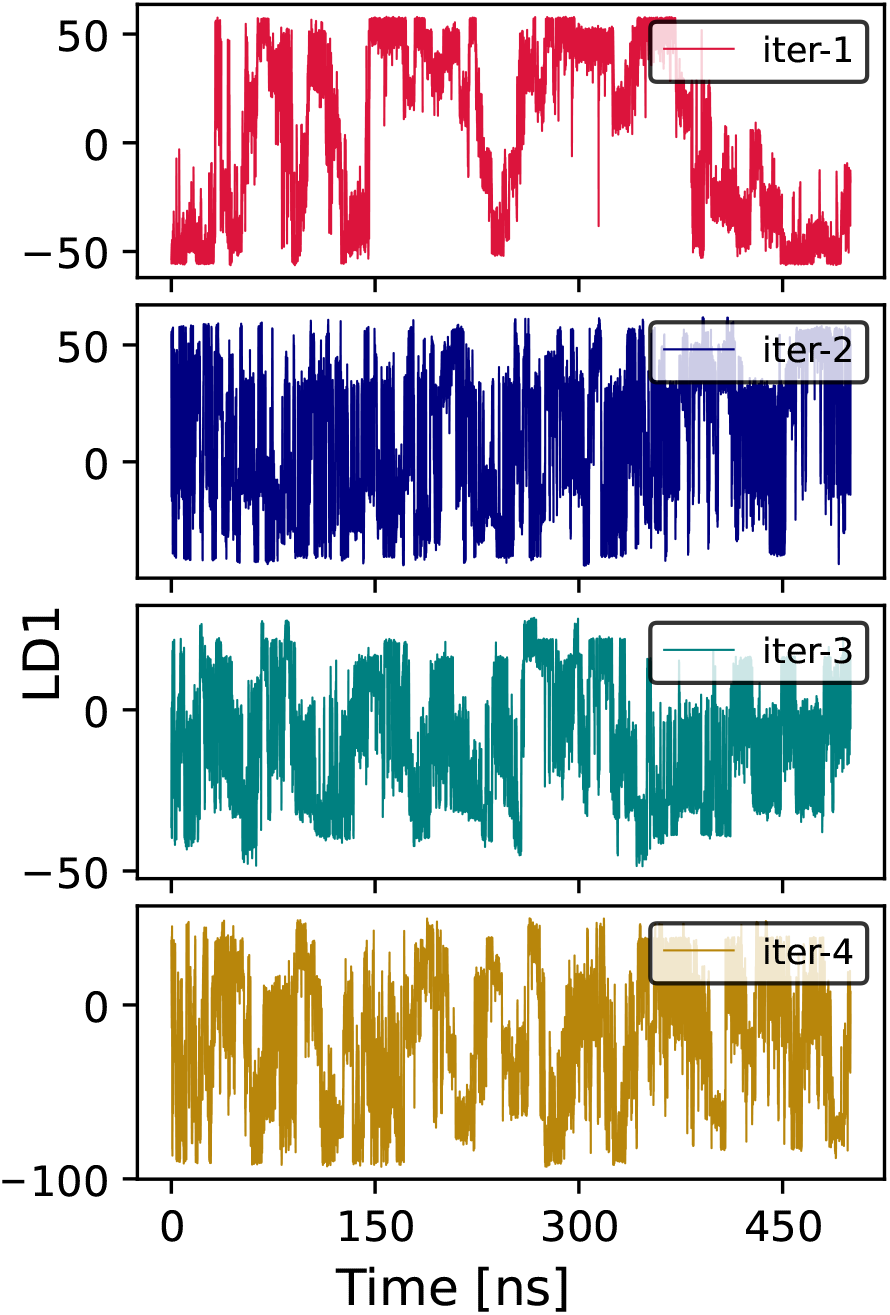
(Aib)_9_ iteration results. Time course of biased LDA coordinate. 500 ns of data were used for clustering and training of the next LD coordinate.

To perform iterations, we scan over possible numbers of clusters, and compute the log-likelihood for each clustering as shown in Fig. S1. These data were used to pick the “best” number of clusters as described in Ref. 26. Fig. S2 displays the calculated Bhattacharyya distances for all newly identified clusters relative to the initial states, for the last three iterations (see Sec. A for details). The two states nearest to either the left or right side are selected to construct a new weighted LD1 coordinate for the subsequent iteration. The magnitudes of particle displacement vectors acting on individual atoms for all LD1 coordinates are depicted in Fig. S3.

From these data, we are also able to compute FE profiles along each LD coordinate, as shown in Fig. S4. Because the coordinate is changing each time, and we do not have a long unbiased reference data set to check convergence, we therefore focus on a previously defined helicity coordinate, ζ, that is the sum of the five central ϕ dihedral angles. ^15^ Values of ζ ∼ − 5 and ζ ∼ +5 correspond to the right- and left-handed helices, respectively. This CV serves as a consistent reference coordinate to track state transitions and assess convergence of the FES along it (Fig. 3 ).

**Figure 3.**
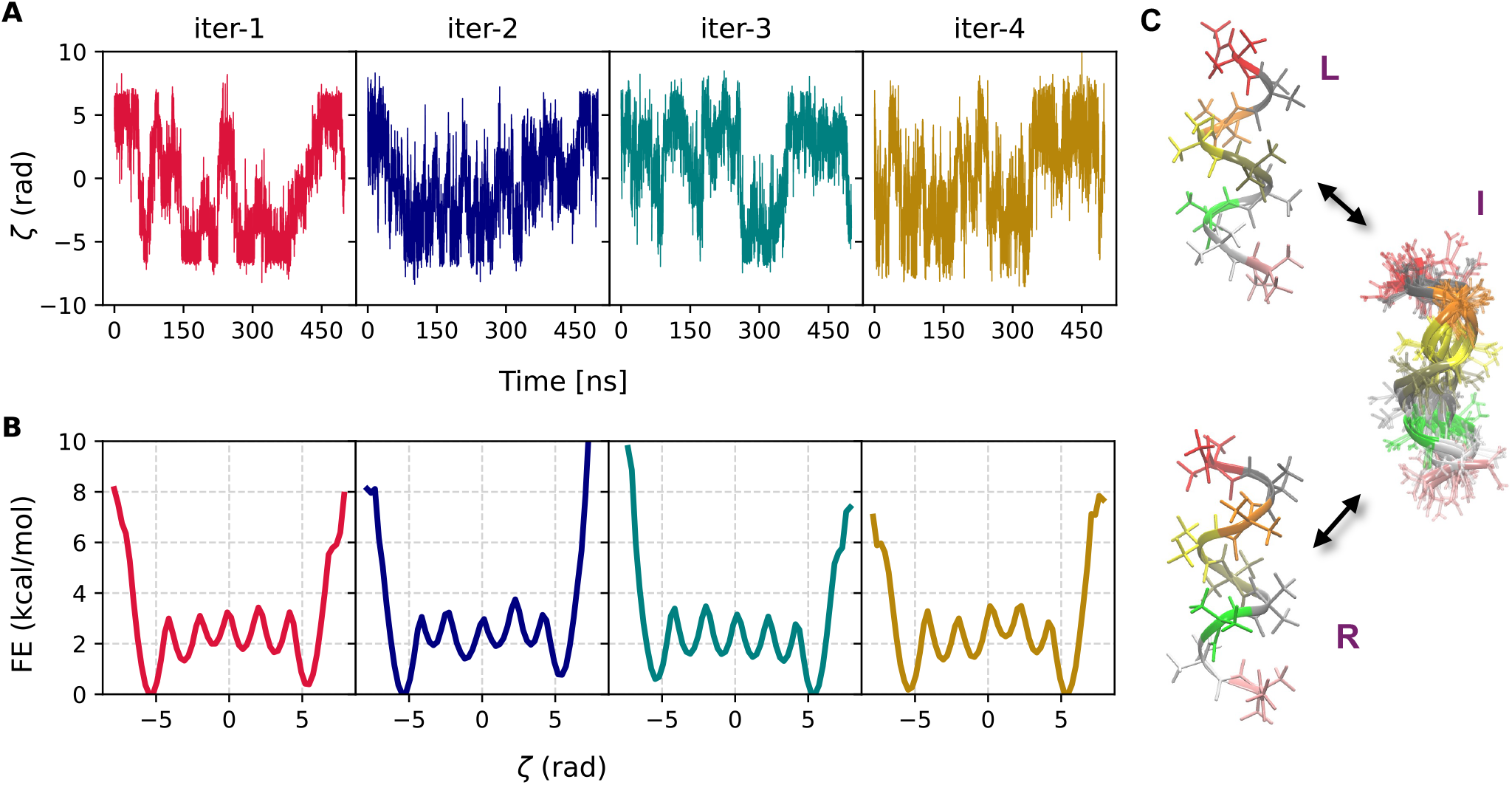
(Aib)_9_ iteration results. (**A**) Trajectory of the physically motivated ζ with time for successive iterations. (**B**) Free energy vs. ζ for successive iterations estimated after 500 ns of MD, and that same data was used to perform the iteration. Each iteration shows transitions between left and right handed states, but iteration four shows the most transitions per unit time and the most symmetric free energy profile from the limited sampling. (**C**) Representative structures from iteration 1, showing transitions from left-(L) to right-(R) helical states of (Aib)_9_. Intermediate (I) shows representative snapshots of configurations having ζ ≈ 0.

Fig. 3 illustrates the transitions along ζ during WT-MetaD simulations across all four iterations, along with the corresponding one-dimensional reweighted free energy profiles. The first coordinate is highly sensitive to the application of bias forces, as previously discussed in Ref. 9, meaning that gentle biasing had to be applied to prevent “crashing” due to rapid changes in forces. This resulted in relatively slow sampling of the configurational space. In contrast, here we observed that the CVs obtained in subsequent iterations are substantially more stable and effective in facilitating extensive sampling of the free energy surface (FES) when employed with higher hill heights and bias factors. For the specific values of the MetaD parameters used, refer to Sec. 5. Notably, state transitions increase significantly from the second to the fourth iteration with consistent MetaD parameters.

To test whether free energy profiles along each coordinate would eventually converge, we extended each simulation to 1.5µs (Fig. S5). Fig. S6 displays ζ fluctuations over time and show that in this time period, all CVs exhibit a FE profile with left and right states having equal free energy minima within 0.5 kcal/mol.

As a final experiment on this system, we explored the effect of enforcing equal contributions from the two states when constructing the weighted LD1 coordinate (i.e., assigning each state a combined sample weight of one). This approach was tested for iteration 2 using biased data from the first enhanced sampling simulation. The resulting coordinate performs comparably to the original, as shown in Fig. S7. These findings demonstrate that iterative refinement, utilizing enhanced sampling data from each step, systematically improves reaction coordinates for (Aib)_9_, enhancing the exploration of its conformational landscape.

### 3.2 Performing iterations on HP35

We previously applied our shapeGMM clustering approach on a 305 µs long MD trajectory of the fast-folding Nle/Nle mutant of HP35, obtained from the D.E. Shaw Research Group. For our analysis, we selected a six-state representation of the system, which provides an interpretable depiction of the folding and unfolding process. Details of the clustering methodology and cross-validation are discussed in Ref. 26. The six-state model was trained using 25,000 frames sampled from a dataset of approximately 1.5 million frames. In our subsequent study, we demonstrated that a single folding/unfolding coordinate could be derived by performing LDA on frames assigned to the folded and unfolded states from this six-state representation. ^9^ Remarkably, this coordinate—trained exclusively on two states—was sufficient to characterize transitions between the folded and unfolded states through physically meaningful configurations. Moreover, it proved to be an effective sampling coordinate when biased in OPES-MetaD simulations ^9^

Here, we have implemented the proposed iteration scheme on the system to assess the effectiveness of our approach. The first iteration aligns with the methodology employed in our prior work. Following the procedure outlined in Section 2.1, we conducted a total of three iterations. In the second and third iterations only 2.5 µs, 1.5 µs of biased data from the previous runs, respectively, were used to train the new wLDA coordinates. The resulting cluster scan for each training iteration are illustrated in Fig. S8. For the second iteration, the training process utilized 44,000 samples, with an additional ∼5,000 samples reserved for cross-validation. In the final iteration, the dataset was expanded to 90,000 training samples, supplemented by 10,000 samples for cross-validation. In each iteration, we computed the Bhattacharyya distance between the newly generated clusters and our predefined folded and unfolded states (see Sec. A). The resulting *D*_*B*_ data, presented in Fig. S9, quantifies the similarity between the two clusters.

The clusters most closely resembling either the folded or unfolded states are selected. The weighted LDA coordinates derived between the new states at each iteration differ from one another, and the coefficients of the LD1 vectors are illustrated in Fig. S10. The variation of the LD1 coordinates from OPES-MetaD simulations, performed for every iteration and the corresponding free energy (FE) profiles computed along them are displayed in Fig. S11. Additionally, Fig. S12 shows the reweighted 2D free energy surface (FES) projected onto the RMSD space, calculated using those biased simulations.

To assess free energy convergence, each OPES-MetaD simulation was extended, and Fig. 4 presents the time dependence of LD1 coordinates and 1D FE profiles obtained along them from the extended simulations for all iterations. While the first simulation ran for approximately 10 µs, we achieved significant convergence to the reference free energy (FE) in the last two iterations within just 3.5 µs. This suggests a notable improvement in the effectiveness of the new coordinates used in iterations 2 and 3. To further evaluate this improvement, we computed the 2D free energy surfaces (FES) projected along the RMSD coordinates relative to Helices 1 and 3 for both simulations, as shown in Fig. 5. The results clearly demonstrate that the new wLDA coordinates served as a more efficient sampling coordinate. The system was able to explore a broader region of the free energy landscape, showing strong agreement with the reference FES derived from D.E. Shaw Research data (depicted as contour lines).

**Figure 4.**
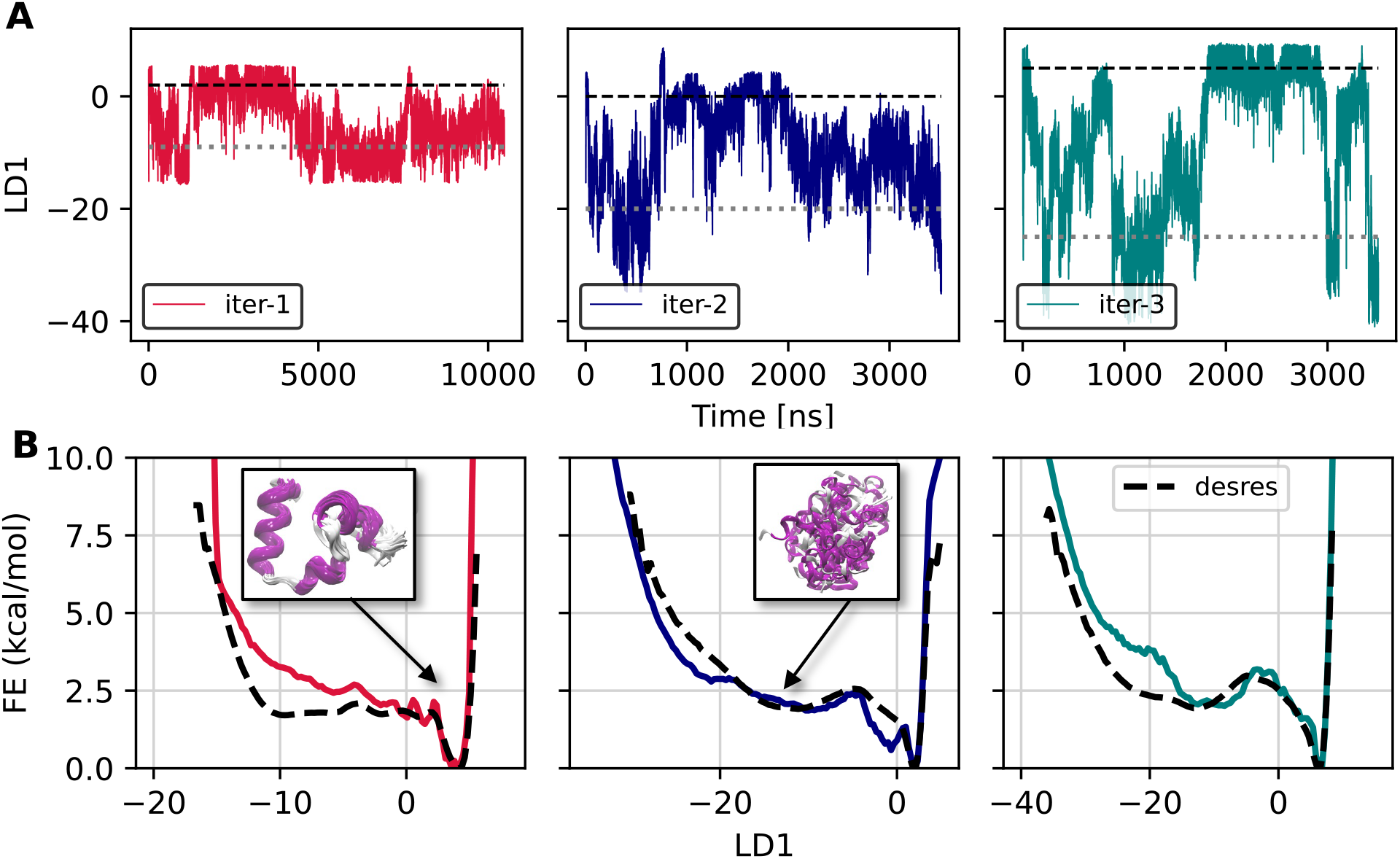
(**A**) Fluctuations of LD coordinates with time from extended OPES-MetaD runs for HP35. Horizontal black dashed and gray dotted lines in each case indicate the approximate locations for the folded and unfolded states respectively. The simulations for iteration 1 ran for 10 µs and the remaining two ran for approximately 3.5 µs. (**B**) FE as a function of LD1 for successive iterations. The black dashed line represents the unbiased free energy estimate derived from D.E. Shaw Research data. ^42^ The improved alignment in iteration 2 and 3 indicates better convergence and enhanced sampling efficiency. The insets highlight representative structures of the folded and unfolded clusters (both taken from iteration 2), illustrating the conformational changes during the transition. Protein conformations are colored according to their secondary structure.

**Figure 5.**
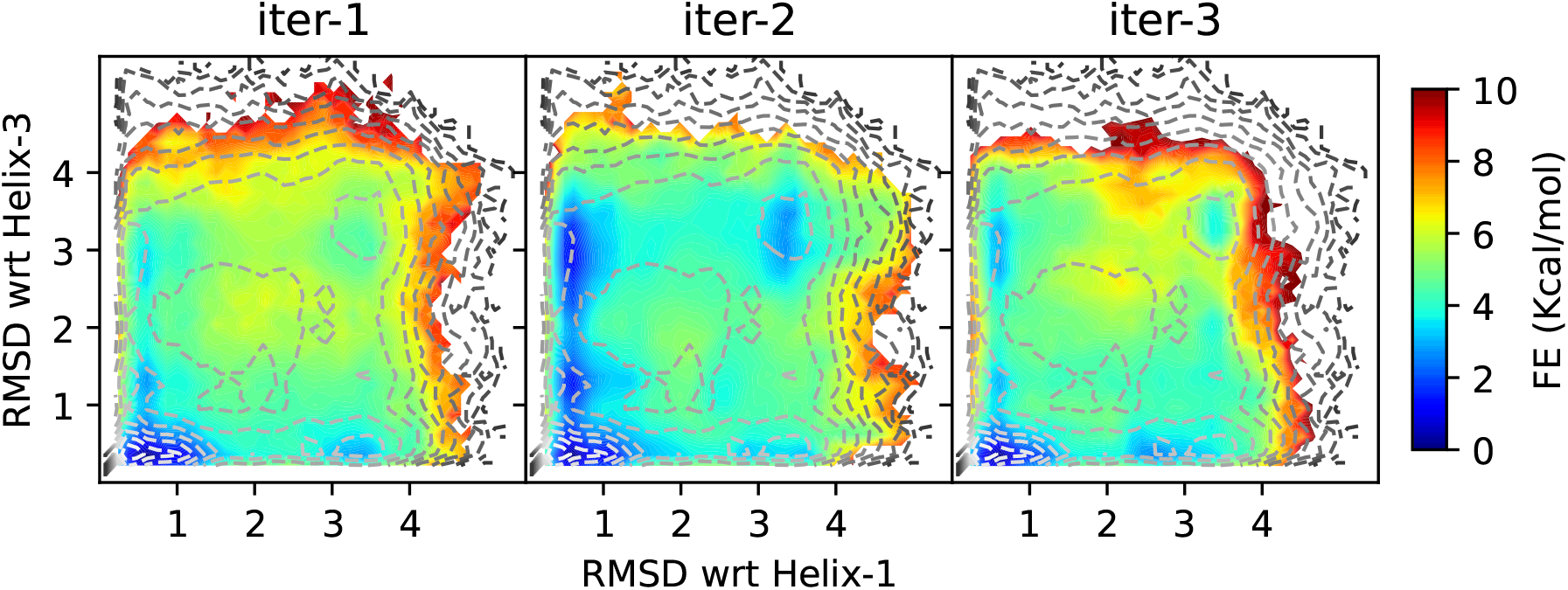
Comparison of 2D FES from three iterations, projected along the RMSD with respect to Helices 1 and 3. The color scale represents free energy in kcal/mol. Contour lines indicate the reference free energy estimate derived from D.E. Shaw Research data. ^42^ The FESs from iteration 2 and 3 demonstrate improved sampling, capturing a broader region of the free energy landscape with better agreement to the reference.

## 4 Conclusions and Outlook

Our results demonstrate that LDA coordinates based on atomic positions can be iteratively improved using data generated by Metadynamics-like sampling. Iteration improves both the stability and sampling efficiency of the resulting CV. We note that these two effects are coupled, in that improved stability of the CV also allows us to bias more quickly, which improves sampling efficiency. Yet in our studies, we also find this is not the only effect, and the CV also allows better exploration even when using a similarly gentle MetaD bias to that from earlier iterations.

We speculate that this results from a better estimate of the positional covariance matrix around each metastable state, which produces a more smooth transition between the two targeted states, however this requires further investigation.

Although our results are promising, some challenges and questions remain. One question in any such approach is: how long should each stage of the iteration be? If a stage of the iteration were run exhaustively to convergence, then there would be no need to iterate to produce a more efficient coordinate. Our results show for these examples that it is possible to improve the coordinate by using enough sampling time to have one to two round trip visits to each state. However, it remains to be investigated whether that generalizes to more challenging systems, and also whether it produces better results than running using the first CV for as long as all of the iterations combined. We believe that there is an actual improvement in CV quality that results in better estimates, for example in Fig. 5, where the target unfolded state at the top right is barely populated even after 10 microseconds of sampling for iteration one, but is properly given weight in a much shorter simulation on the second iteration.

Going forward, we would like to build on this approach for sampling more challenging systems. When going to a larger system, we believe based on our experience that a single linear coordinate will not be sufficient to capture all the slow degrees of freedom when transitioning between two states. We therefore would like to investigate combining iterative improvement of one posLDA CV that separates the two states of interest, with other CVs that promote exploration of other large domain motions. We also would like to investigate cases that include metastable intermediates, to see whether one single posLDA between the end states is a good CV for sampling, or whether we should in fact combine multiple posLDA CVs defined pairwise between metastable states, which would allow us to better sample from e.g. a starting state to an intermediate and then from the intermediate to a final state via a more physically realistic route.

## 5 Simulation Details

All simulations were performed using GROMACS 2020.4^43^ with PLUMED 2.9.0-dev. ^25,36^ All analysis scripts, Jupyter-notebooks and PLUMED input files used in the study are currently available in our paper’s GitHub repository https://github.com/hocky-research-group/Sasmal_posLDA_iteration, and will also be available on Zenodo and PLUMED-Nest ^25^ on publication.

### (Aib)_9_ Simulations

Equilibrated inputs for (Aib)_9_ were provided by the authors of Ref. 15. In brief, simulations using the CHARMM36m forcefield and TIP3P water. ^44^ MD simulations are performed in NPT with a 2 fs timestep at *T* = 400*K*. The MetaD parameters used for first iteration were HEIGHT=0.005, BIASFACTOR=2, SIGMA=0.43, PACE=500. For all three remaining iterations we used, HEIGHT=0.70, BIASFACTOR=8, SIGMA=0.55 and PACE=500 and a multiple time step STRIDE for biasing of 2. ^45^ Quadratic upper and lower walls were applied ∼ ±10.0 of maximum and minimum value for each LD1 coordinate respectively, with a bias coefficient of 125 kcal/mol/Å^2^. Complete details are provided in `PLUMED` input files on GitHub.

### HP35 Simulations

A 305 µs all-atom simulation of Nle/Nle HP35 at *T* = 360*K* from Piana et al. ^42^ was analyzed. The simulation was performed using the Amber ff99SB*-ILDN force field and TIP3P water model. In that simulation, protein configurations were saved every 200 ps, for a total of ∼1.5M frames. For our simulations, we solvate and equilibrate a fresh system using the same forcefield at 40mM NaCl. Minimization and equilibration are performed using a standard protocol^1^, at which point NPT simulations are initiated at *T* = 360*K*. mdp files for all steps of this procedure and the topology files are all available in the GitHub of our previous work. ^9^ All the OPES-MetaD simulations are performed with γ = 8, Δ*E* = 10 kcal/mol, pace of 500 steps, and a biasing multiple time step ^45^ stride of 2. Quadratic walls were applied for each LD1 coordinate, specific to its range between upper and lower limits, with a bias coefficient of 125 kcal/mol/Å^2^.

### A Bhattachrayya Distance

The Bhattacharayya distance is a statistical measure used to quantify the similarity between two probability distributions. It is derived from the Bhattacharyya coefficient which measures the amount of overlap between two distributions. For any two given continuous distributions *p*(*x*) and *q*(*x*), the coefficient is defines as

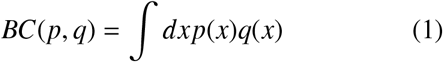

and the distance *D*_*B*_ is given by,

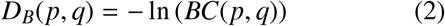

*D*_*B*_ is a symmetric quantity that means *D*_*B*_(*p, q*) = *D*_*B*_(*q, p*). It goes to zero for identical distributions and goes to infinity for entirely dissimilar distributions. It is assumed that the compared distributions are well defined and normalized. If the distributions are too different from each other, the distance can be very large.

If *p*(*x*), *q*(*x*) are multivariate normal distributions such as *p*(*x*) ∼ 𝒩 (µ_*p*_, Σ_*p*_) and *q*(*x*) ∼ 𝒩 (µ_*q*_, Σ_*q*_), then it can be derived to show that *D*_*B*_(*p, q*) is given as,

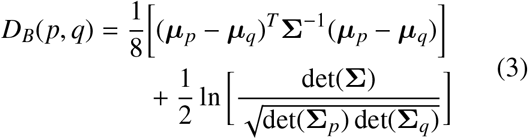

***µ***_*p*_, ***µ***_*q*_ are the mean vectors corresponding to distributions *p*(*x*) and *q*(*x*) with covariances **Σ**_*p*_, **Σ**_*q*_ respectively. And 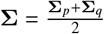 , is the mean of two covariances. The first term in the Eq.3 is the Mahalanobis distance between two distributions that quantifies the difference in their locations. The second term accounts for the difference in the shapes of the two distributions and is a measure of the divergence due to differences in the spreads and orientations.

For this work, we implemented the Bhattacharayya distance metric in the similarities module of the shapeGMMTorch package. ^24^

## Acknowledgement

We thank the D. E. Shaw Research for providing simulation data on the HP35 protein and we thank the Tiwary lab for providing their input files for (Aib)_9_. SS and GMH were supported by the National Institutes of Health through the award R35GM138312. SS was also partially supported by a graduate fellowship from the Simons Center for Computational Physical Chemistry (SCCPC) at NYU (SF Grant No. 839534). MM would like to acknowledge funding from National Institute of Allergy and Infectious Diseases of the National Institutes of Health under award number R01AI166050 and the National Science Foundation under award 2238706. MM would like to acknowledge Peter T. Lake, who implemented the frame-weighted LDA method as a Python package. ^35^ This work was supported in part through the NYU IT High Performance Computing resources, services, and staff expertise, and simulations were partially executed on resources supported by the SCCPC at NYU.

## Supporting information

### S1 Training curves for (Aib)_9_ iterations

**Figure S1:**
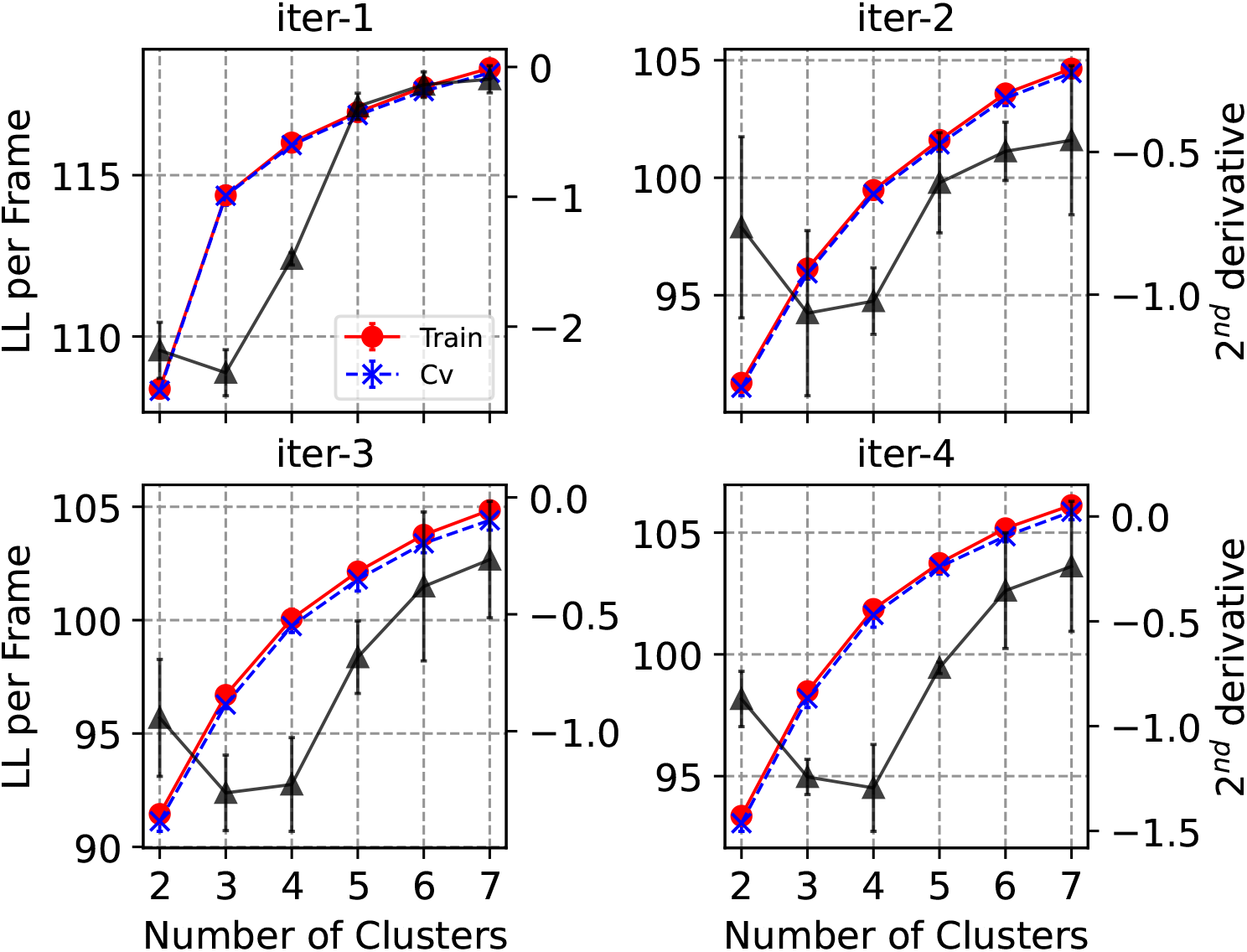
Cluster scans from four successive iterations of (Aib)_9_. Iteration 2,3 and 4 were performed with data from biased simulations, using 90k training samples and 10k samples for cross validation. First iteration was performed with combined data from two short 20ns long MD simulations initiated form both left and right states. In first iteration, we used 20k frames for training along with 20k for cross validation. Black curves represent 2^*nd*^ derivatives(with error bars) of log likelihood with respect to number of clusters and minimum value indicates an optimal choice for number of clusters.

### S2 Bhattacharyya Distances for (Aib)_9_

**Figure S2:**
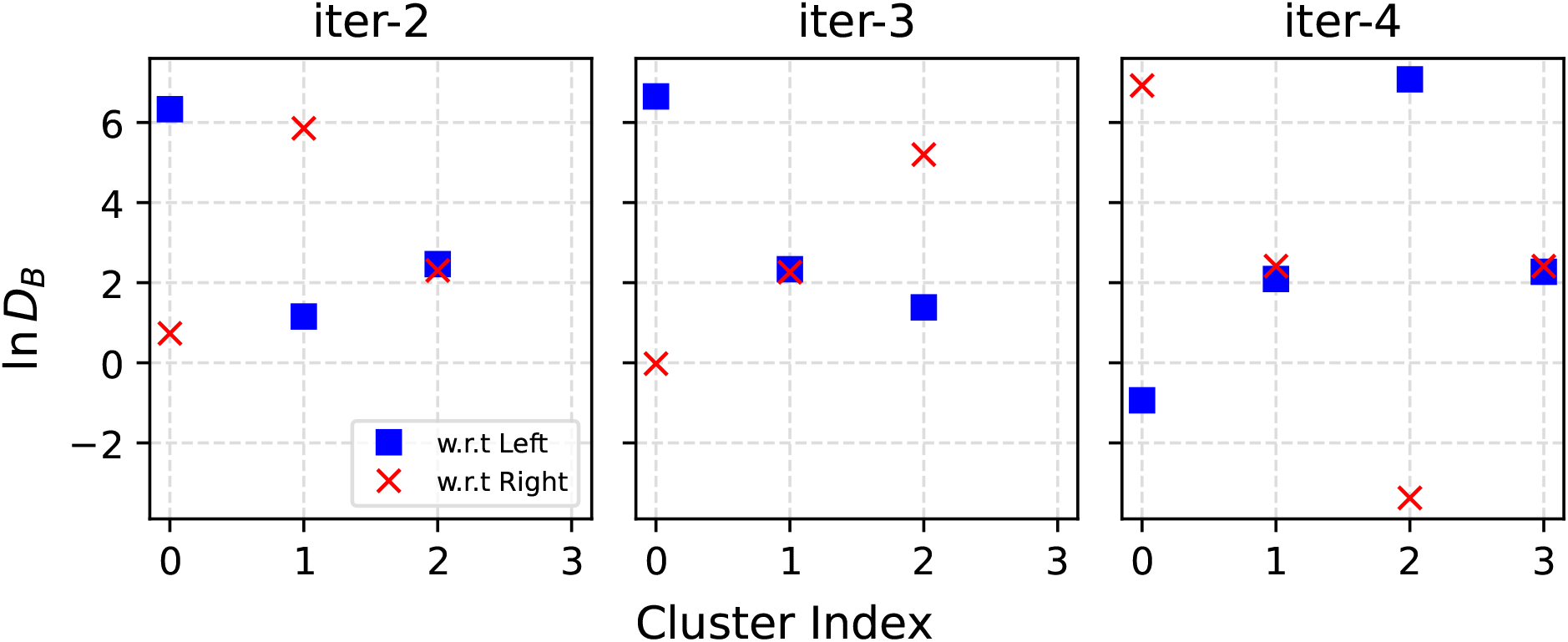
Logarithm of Bhattacharyya distances for all clusters with respect to initial definitions of left and right helical states at every iteration. It gives a measure of similarity between two multivariate normal distributions that represent a cluster. Any two clusters with lower values of ln *D*_*B*_ are close to each other and those with higher values are far away from each other. It provides a consistent way of defining new left and right states at every iteration in accordance with initial definitions.

### S3 Coefficients of LD coordinates from (Aib)_9_ iterations

**Figure S3:**
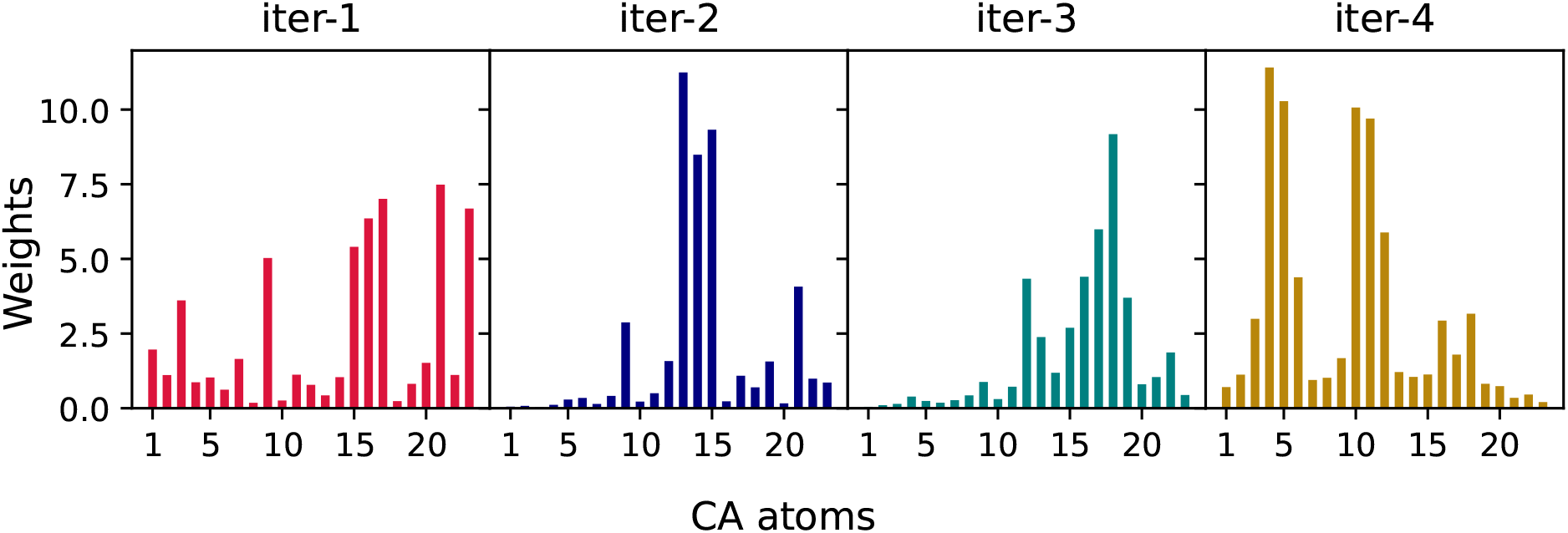
Weights shown are the magnitudes of particle displacement vectors acting on each atom from LD1 after each iteration. In case of (Aib)_9_, cartesian coordinates of total 23 backbone atoms are used to define LD1 coordinate which is a linear combination of 23 × 3 = 69 features with 69 real coefficients. Hence, each particle has a displacement vector of 3 components associated with it. In this figure, it shows the magnitude of those vectors. Weights are considered as contributions of different atoms in making the coordinate. The atoms with larger weights have a larger effect when biasing while those with smaller weights contribute less.

### S4 FEs vs. LD1 for (Aib)_9_ iterations

**Figure S4:**
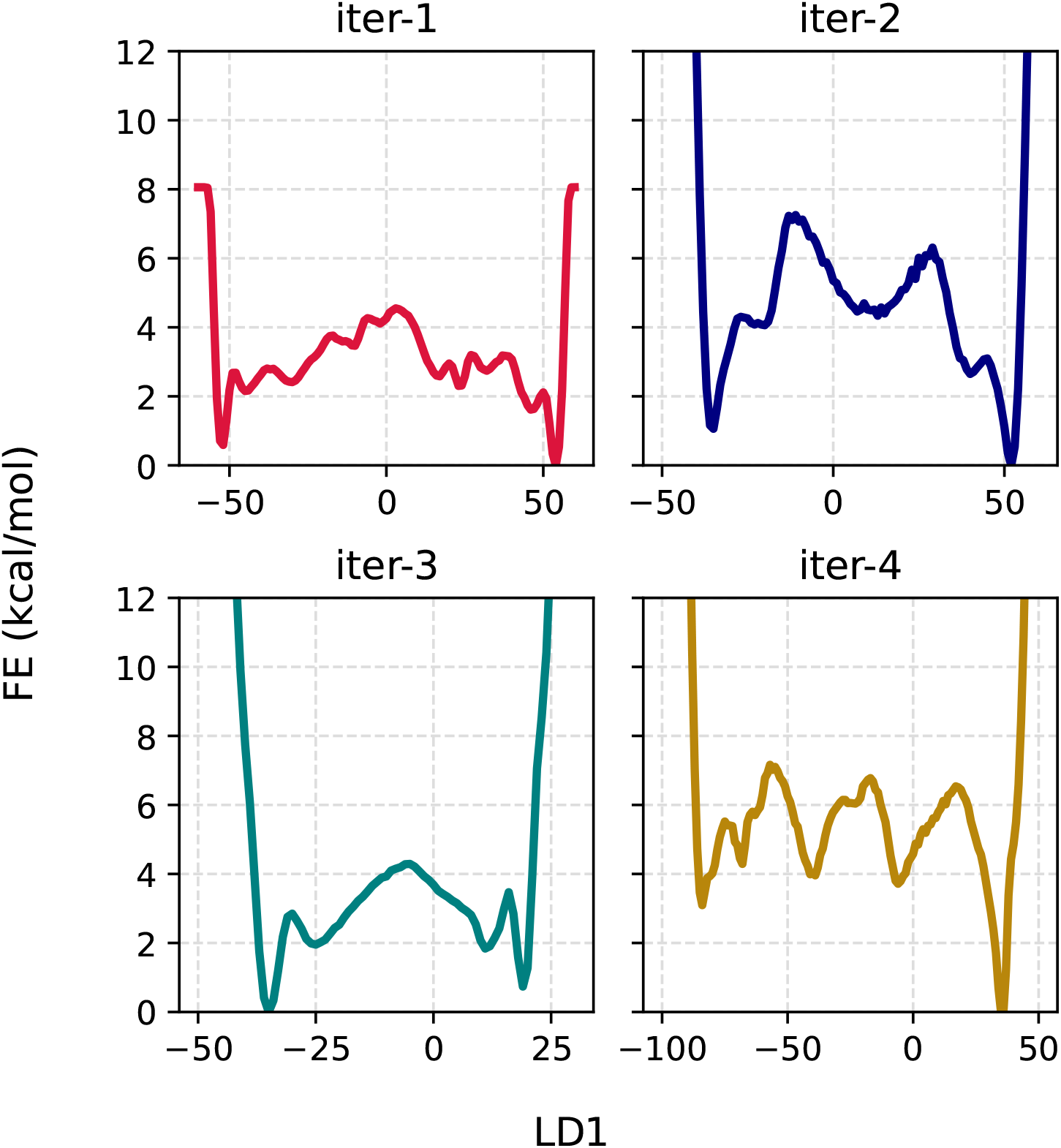
FE profiles along LD1 obtained from 500ns long WT-MetaD simulations in four successive iterations of (Aib)_9_.

### S5 FEs and time dependence of LD coordinates from (Aib)_9_ 1.5*μ*s simulations

**Figure S5:**
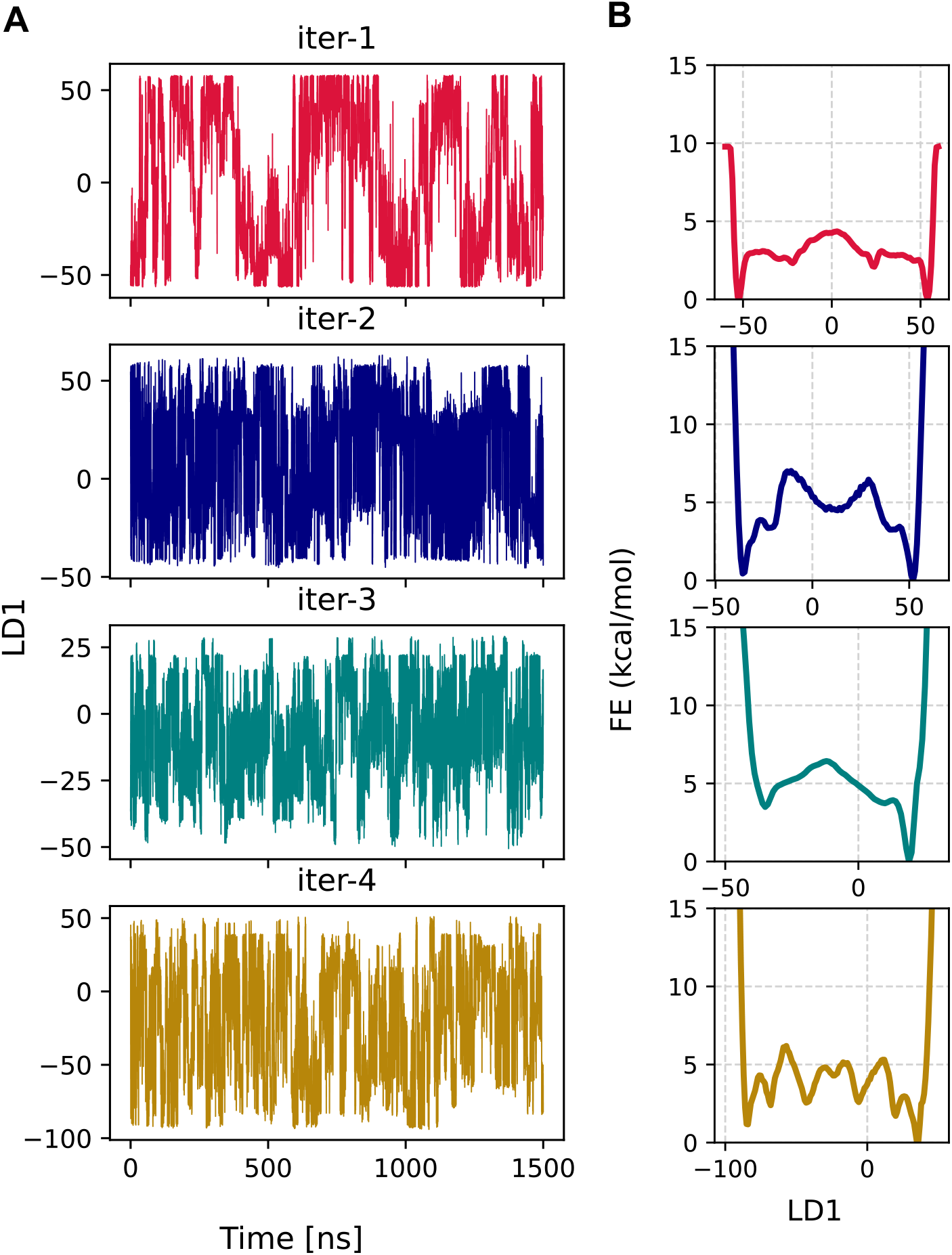
Each WT-MetaD simulation from successive iterations were further extended upto 1.5 µs. (**A**) Fluctuations of LD1 with time and (**B**) FE profiles computed along LD1 in each case by summing all Gaussian hills deposited over the course of the simulations.

### S6 FEs and time dependence of *ζ* from (Aib)9 1.5*μ*s simulations

**Figure S6:**
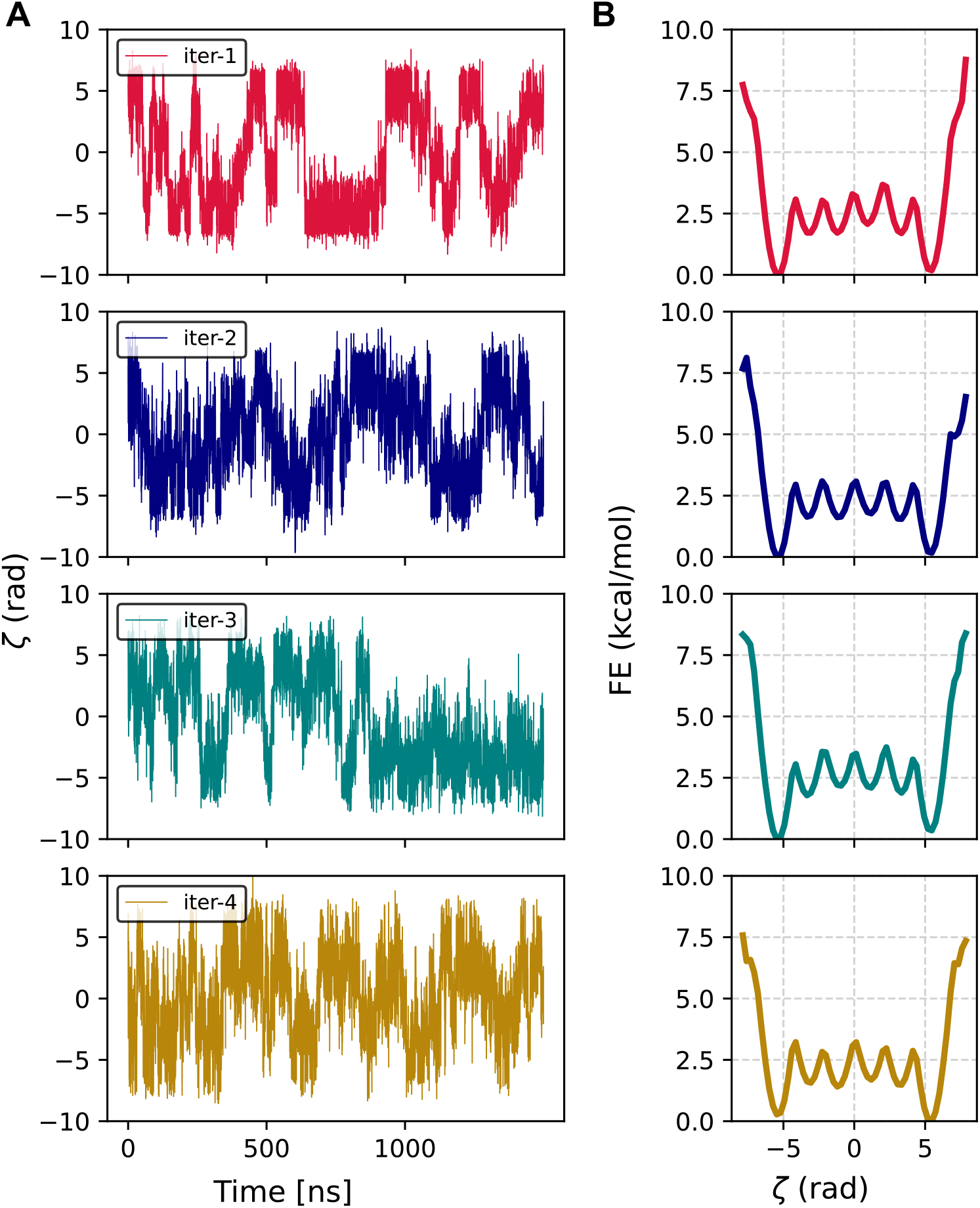
(**A**) fluctuations of ζ with time and (**B**) converged reweighted free energy profiles computed from 1.5µs long WT-Metad simulations. Efficient sampling between left and right states is observed in all cases. All free energy profiles along ζ are converged and symmetric.

### S7 Implementing Equal Weights for left and right helix

**Figure S7:**
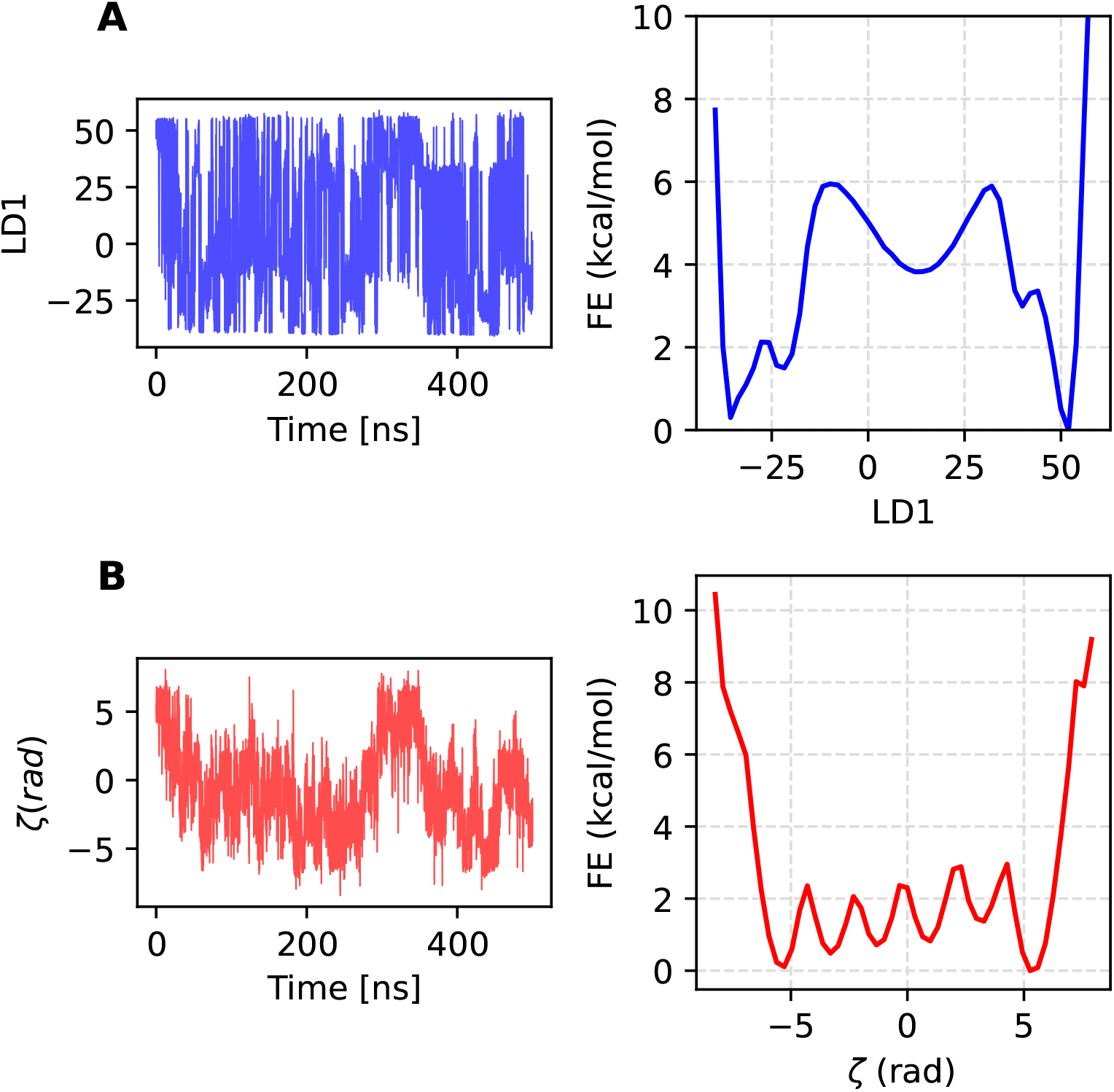
Results from a 500ns long WT-MetaD simulation by biasing LD1 coordinate which is obtained by implementing equal total probability for samples belonging to left and right states. Equal total probability means that sum of weights for all the samples from either left or right state is equal to 1. To test this we used WT-MetaD data from first iteration. After computing the correct weights for samples from biased data, here we normalized the weights separately for left and right states before feeding it into LDA algorithm, so that each state contributes equally to the coordinate. (**A**) LD1 vs. time and free energy profile calculated along LD1 and (**B**) ζ vs. time along with FES along ζ. The MetaD parameters used for this simulation are, HEIGHT=0.01, BF=8, PACE=2000, SIGMA=0.55 and STRIDE=2. Two quadratic walls were applied at LD1=+60.0 and LD1=-60.0 with force constant of 125.0 kcal/mol/Å^2^.

### S8 Training curves for HP35 iterations

**Figure S8:**
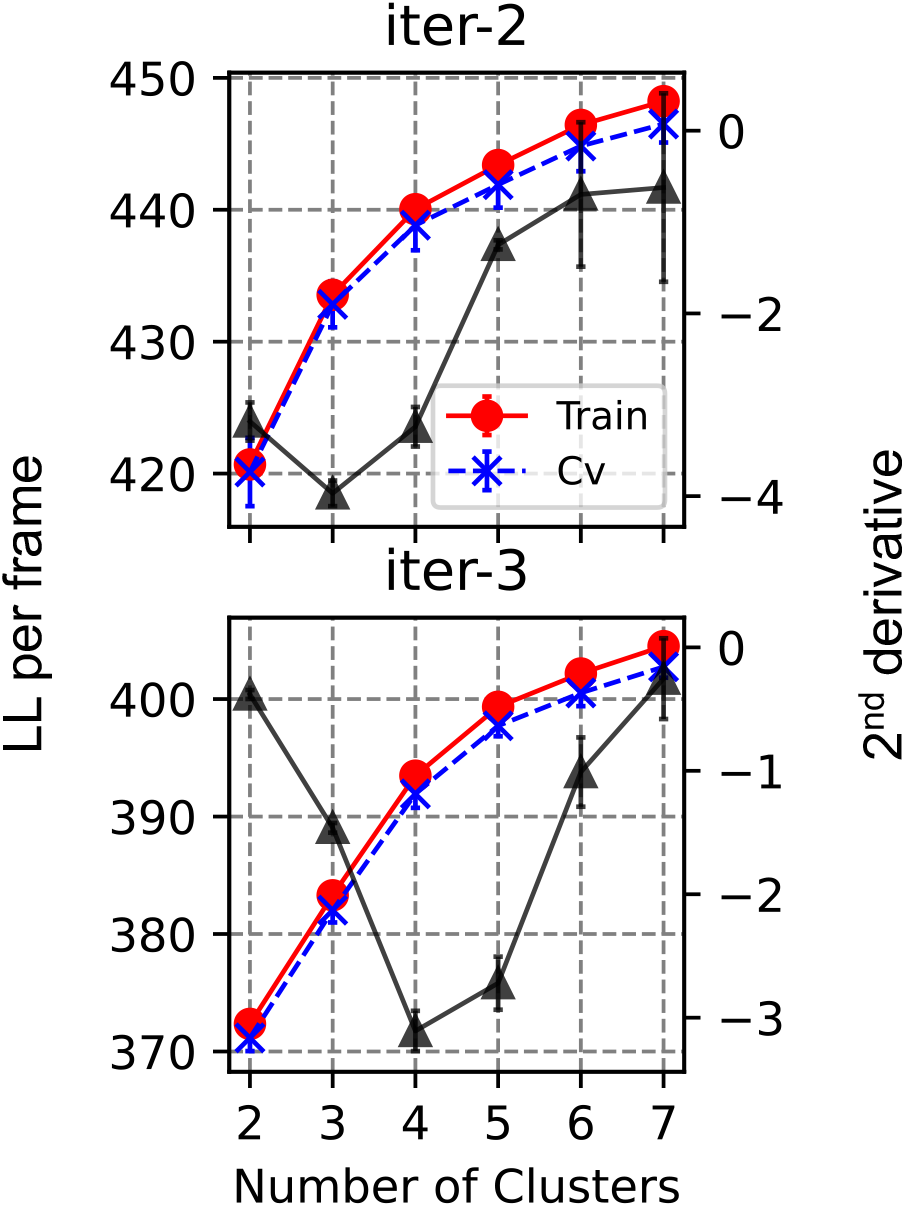
Cluster scans for last two iterations of HP35. The first scan (reported in Ref. 1, not shown here) was performed with 305µs long MD simulation trajectory of HP35 provided by D. E. Shaw Research. ^2^ The second scan was performed with 2.5µs long OPES-MetaD simulation data with 44k frames for training along with ∼5k frames for cross validation. The third scan was performed with 90k samples for training and 10k samples for cross validation, using the biased data from previous 1.5µs long OPES-MetaD simulations. Training curves with error bars are shown in red and cross validations curves with error bars are shown in blue. Black curves represent 2^*nd*^ derivatives(with error bars) of log likelihood with respect to number of clusters and minimum value indicates an optimal choice for number of clusters.

### S9 Bhattacharyya Distances for HP35

**Figure S9:**
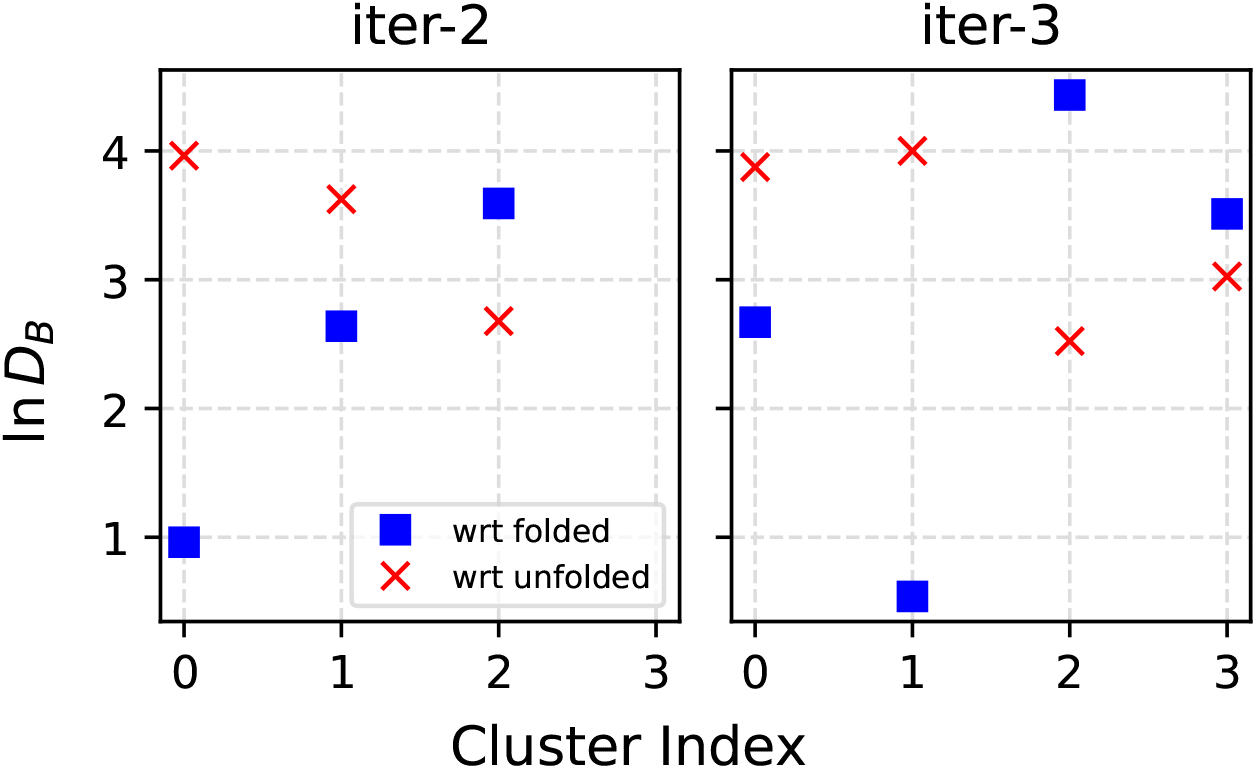
Logarithm of Bhattacharyya distance for all clusters (see Fig. S8) in our HP35 iterations with respect to initial definitions of folded and unfolded clusters.

### S10 Coefficients of LD coordinates from HP35 iterations

**Figure S10:**
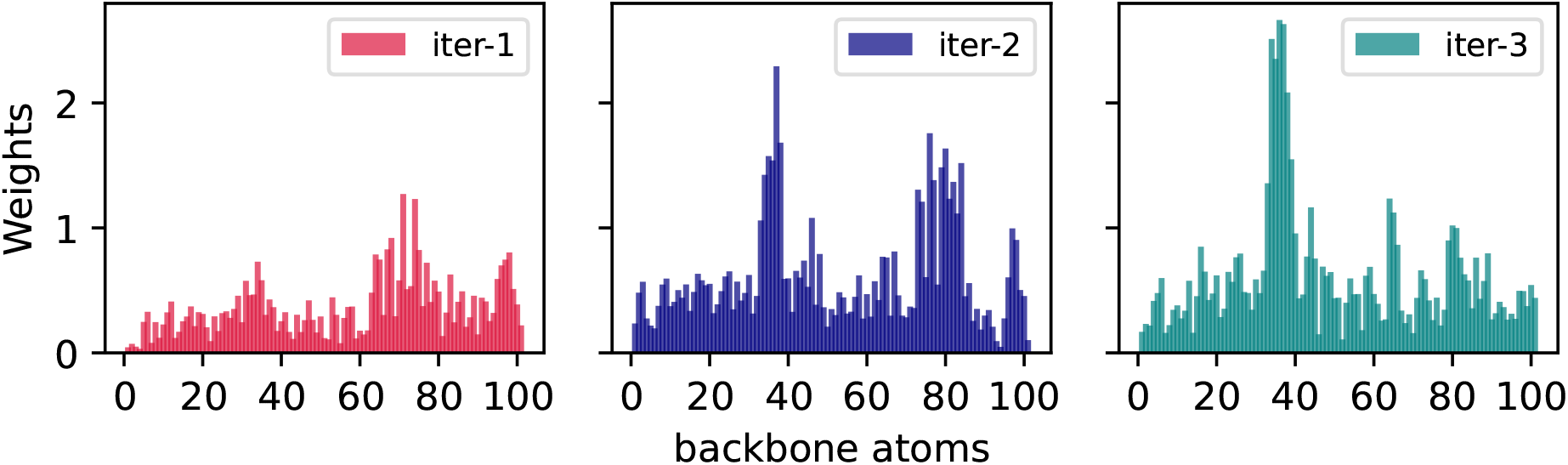
LDA weights at each iteration for HP35. Here, input cartesian coordinates consist of 101 backbone atoms, which is a linear combination of 101 × 3 = 303 features with 303 real coefficients.

### S11 FEs vs. LD1 for HP35 iterations

**Figure S11:**
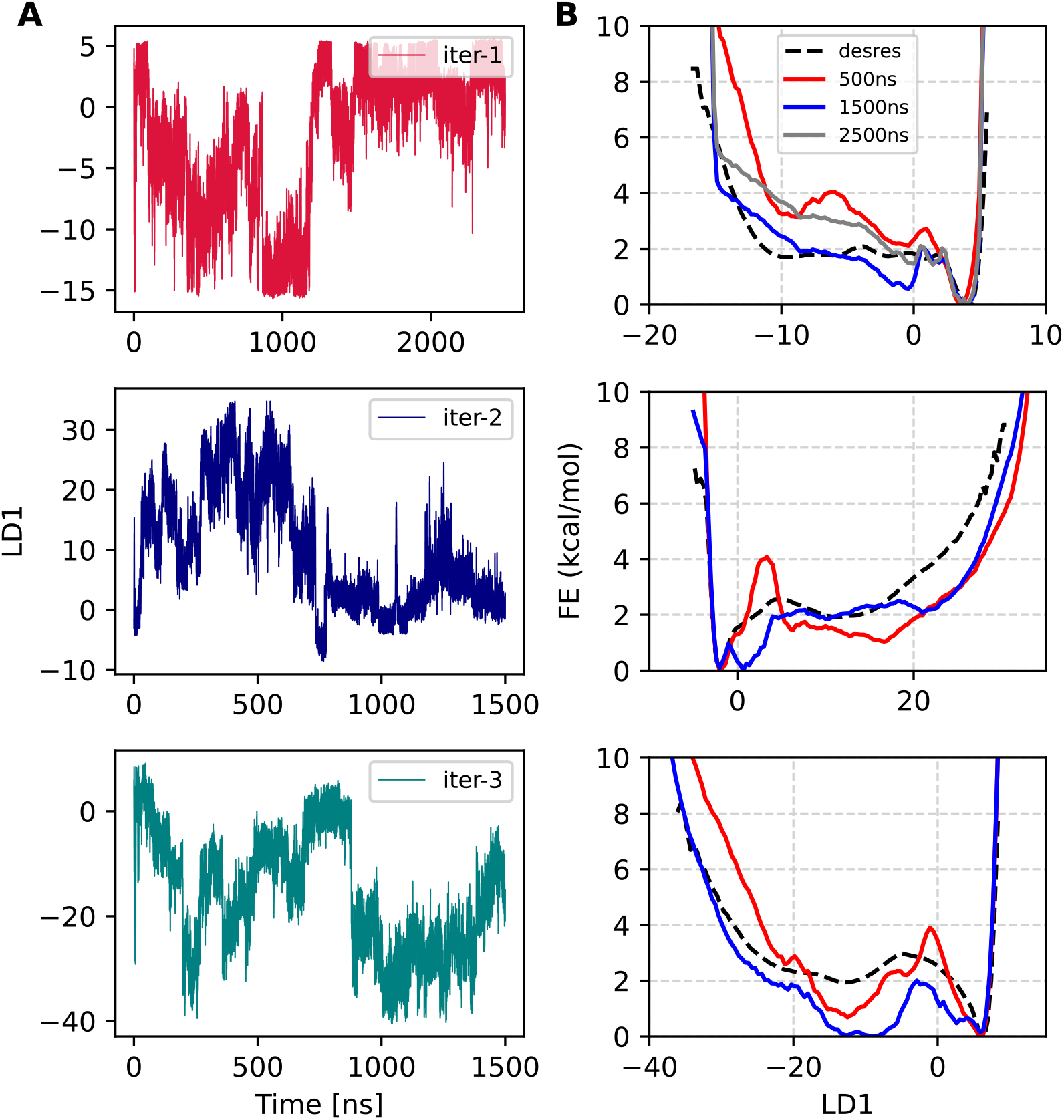
(**A**) Trajectory of LD1 obtained from OPES-MetaD simulations in three successive iterations of HP35. The first simulation is 2.5µs and the remaining two are 1.5µs long. Note that the coordinate obtained at each iteration is different than others. (**B**) FE profiles computed in each oteration. FE profiles calculated using 500ns, 1500ns and 2500ns long data are shown in red, blue, grey colors respectively. In each case for comparison, we also computed a reference FE using 305µs long unbiased MD simulation of villin, provided by D. E. Shaw Research. ^2^ The reference FE profiles are shown in black dashed lines.

### S12 2D FES on RMSD space from HP35 iterations

**Figure S12:**
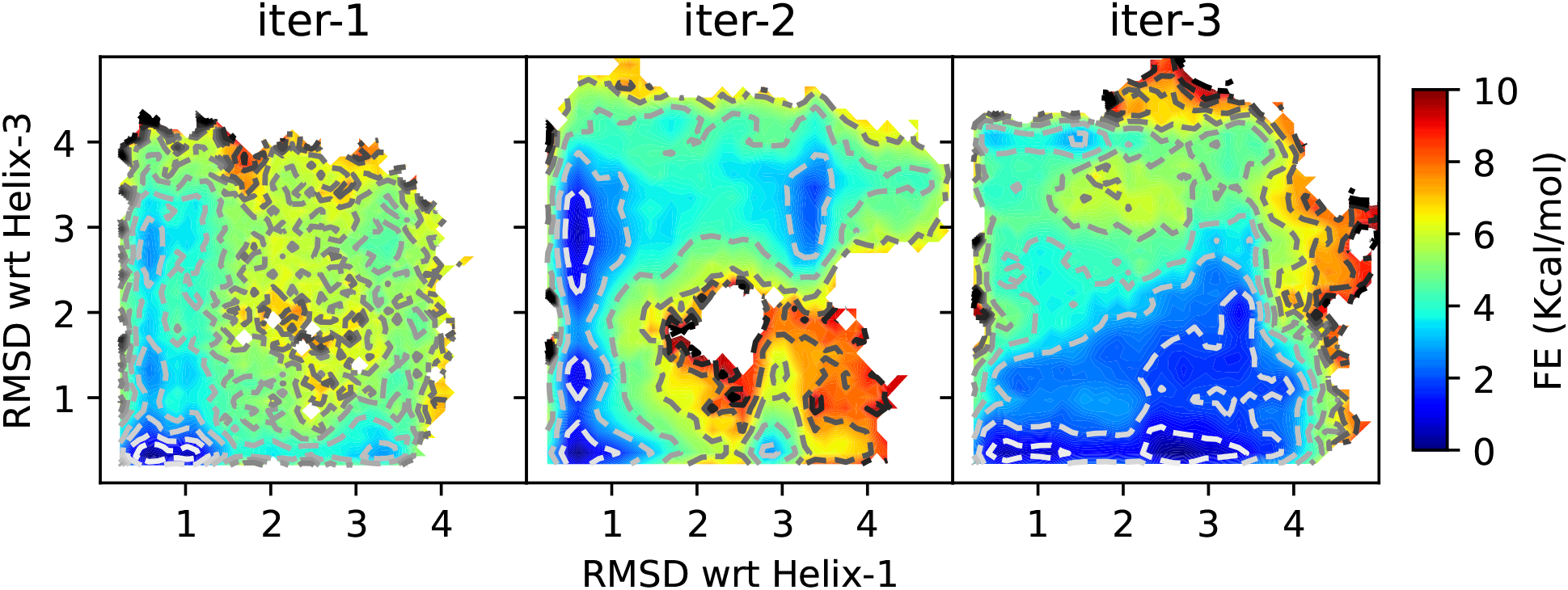
2D reweighted FES projected along RMSDs (computed using only backbone atoms) with respect to helix-1 and helix-3. To compute the free energy profiles, OPES-MetaD simulation data generated at each iteration is used. The first one is 2.5µs long and the later two are 1.5µs long only (see Fig. S11). This also illustrates the input data which is used in an iteration to generate the next wLDA coordinate.

## Footnotes

1 http://www.mdtutorials.com/gmx/lysozyme/index.html

